# Measurements of the timescale and conformational space of AMPA receptor desensitization

**DOI:** 10.1101/847202

**Authors:** Hector Salazar, Sabrina Mischke, Andrew J. R. Plested

## Abstract

Ionotropic glutamate receptors (iGluRs) are ligand gated ion channels that mediate excitatory synaptic transmission in the central nervous system (CNS). Desensitization of the AMPA-subtype following glutamate binding appears critical for brain function, and involves rearrangement of the ligand binding domains (LBDs). Recently, several full-length structures of iGluRs in putative desensitized states were published. These structures indicate movements of the LBDs that might be trapped by cysteine crosslinks and metal bridges. We found that cysteine mutants at the interface between subunits A and C, and lateral zinc bridges (between subunits C & D or A & B) can trap freely-desensitizing receptors in a spectrum of states with different stabilities. Consistent with close approach of subunits during desensitization processes, introduction of bulky amino acids at the A-C interface produced a receptor with slow recovery from desensitization. Further, in wild-type GluA2 receptors, we detected population of stable desensitized state with a lifetime around 1 second. Using mutations that progressively stabilise deep desensitize states (E713T & Y768R), we were able to selectively protect receptors from crosslinks at both the diagonal and lateral interfaces. Ultrafast perfusion enabled us to perform chemical modification in less than 10 ms, reporting movements associated to desensitization on this timescale within LBD dimers in resting receptors. These observations suggest small disruptions of quaternary structure are sufficient for fast desensitization, and that substantial rearrangements likely correspond to stable desensitized states that are adopted relatively slowly, on a timescale much longer than physiological receptor activation.

**Significance statement:** iGluRs are central components of fast synaptic transmission in the brain. iGluR desensitization occurs as a natural consequence of receptor activation and can reduce the response of an excitatory synapse. AMPA receptor desensitization also appears necessary for proper brain development. Molecular structures of iGluRs in putative desensitized states predict a range of movements during desensitization. In the present study, we performed a series of crosslinking experiments on mutant receptors that we subjected to similar desensitizing conditions over time periods from milliseconds to minutes. These experiments allowed us to count desensitized configurations and rank them according to their stabilities. These data show that large-scale rearrangements occur during long glutamate exposures that are probably not seen in healthy brain tissue, whereas smaller changes in structure probably suffice for desensitization at synapses.

## Introduction

Glutamate receptor ion channels mediate most of the fast excitatory synaptic transmission in the vertebrate CNS (Traynelis et al., 2010). Glutamate binding initiates opening of an integral ion pore, permitting cations to flow into the postsynaptic cell. The α-amino-3-hydroxy-5-methyl-4-isoxazolepropionic acid (AMPA) receptor subtype desensitizes rapidly and profoundly in response to the sustained presence of glutamate for more than about 25 ms (Colquhoun et al., 1992; Robert et al., 2005). The number of receptors available to respond to glutamate is consequently reduced during a phase of recovery from desensitization, which in turn can determine the amplitude of postsynaptic responses (Saviane and Silver, 2006; Crowley et al., 2007). The timescale of recovery from desensitization, being for AMPA receptors in the order of tens to hundreds of milliseconds, is pertinent during high frequency release of glutamate (above 10 Hz). Steady-state desensitization may offer protection during pathological glutamate insults that lead to brain damage (Bowie, 2008) and during development (Christie et al., 2010). These observations provide motivation for understanding the molecular basis of AMPA receptor desensitization.

AMPA receptors assemble from four subunits, each comprising an extracellular amino-terminal domain (ATD), a ligand binding domain (LBD) that is connected to the ion channel forming transmembrane domain (TMD) and a carboxyl-terminal domain (CTD). The LBD is formed from an upper D1 and a lower D2 lobe. Upon binding of glutamate, the LBD closes. This motion provokes the separation of the D2 domains leading to the opening of the receptor gate (Armstrong and Gouaux, 2000). Following activation, the receptor transits to desensitized states in about 10 ms, at least in part due to dissociation of the dimer interface formed by the D1 domains (Armstrong et al., 2006).

Recent structures of the full-length AMPA receptor in putative desensitized states suggest a wide conformational space is sampled. Initial cryo-EM structures of the GluA2 receptor in desensitizing conditions showed a set of three-dimensional classes in which the extracellular domains were progressively spread out (Meyerson et al., 2014). A 5-fluorowillardiine-bound structure of GluA2 (Dürr et al., 2014) also showed a large rearrangement of the ATDs and LBDs. Structures of the related GluK2 kainate receptor show the LBDs adopt a 4-fold symmetric arrangement (Meyerson et al., 2016) with individual subunits rotating by more than 120° from their active state dimer positions (Plested et al., 2008) (Figure 1). More recent desensitized state structures of GluA2 in presence of the accessory proteins GSGL1L or TARP γ-2 revealed compact desensitized arrangements, with the LBD dimers losing their internal 2-fold rotational symmetry (Twomey et al., 2017; Chen et al., 2017).

**Figure 1.**
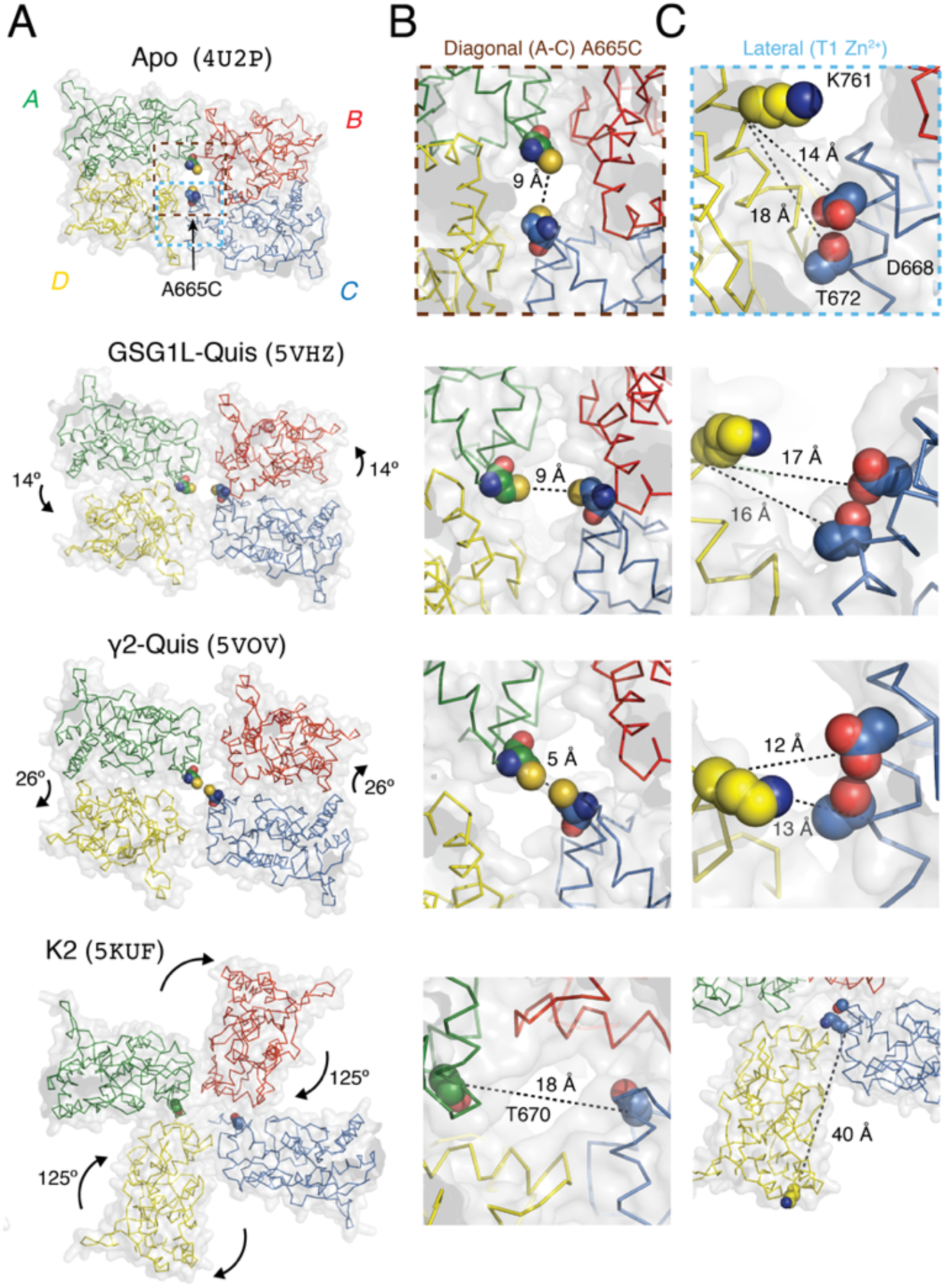
Candidate structures of glutamate receptors in desensitized states. (A) Plan views of the LBD layer in apo (Dürr et al., 2014), 2xGSG1L Quis (Twomey et al., 2017), γ-2 Quis (Chen et al., 2017) and GluK2 (2S,4R-4-methylglutamate; Meyerson et al., 2016) structures. PDB codes shown in brackets. Conformational changes relative to resting/active structures are indicated. Modelled A665C residues are shown as spheres. The scope of insets in panels B and C is shown by brown and light blue dashed boxes, respectively. (B) The A665C mutant site in the A and C subunits. Distances are modelled between Cysteine sulphur atoms, except for GluK2 where the equivalent residue is T670, which is buried. (C) The lateral interface between subunits C&D at the site of the zinc bridge mutant T1. Distances are measured between main chain C-alpha atoms. Nearest match residues for the T1 site in K2 are S669, K673 and K759.

We previously used cysteine and metal-bridge crosslinking to identify compact arrangements of the LBD tetramer associated with activation of the AMPA receptor. These include the “closed angle” conformation (Lau et al., 2013), other compact forms for LBDs fully bound to glutamate (Baranovic et al., 2016) and the partial agonist 5-fluorowillardiine (Salazar et al., 2017). The latter assembly featured a parallel shift of the individual dimers. Bifunctional cysteine crosslinkers on the other hand, indicate that LBD tetramer opens up in both active and desensitized states (Baranovic and Plested, 2018).

In the present study, we revisited our earlier observation that desensitized receptors can be crosslinked very stably between A & C subunits by the A665C disulfide bond (Lau et al., 2013). Using a fast perfusion system, we used several approaches on mutant and wild-type receptors to count conformational states attained during desensitization. Comparing potential desensitized states obtained in crystallographic and cryo-electron microscopy to the geometric constraints imposed by disulfide bonds and metal bridges suggests that compact desensitized arrangements can best account for desensitization on the physiological timescale.

## Materials and methods

### Electrophysiology

All mutants were generated on the GluA2flip background by overlap PCR and confirmed by double-stranded DNA sequencing. For consistency with previous reports, the numbering of mutated amino acids assumes a 21-residue signal peptide for GluA2. Wild type (WT) and mutant AMPA receptors were expressed transiently in HEK-293 cells for outside-out patch recording. All patches were voltage clamped between –30 to –60 mV. Currents were filtered at 1-10 kHz (–3 dB cutoff, 8-pole Bessel) and recorded using Axograph X (Axograph Scientific) *via* an Instrutech ITC-18 interface (HEKA) at 20 kHz sampling rate.

The external solution in all experiments contained 150 mM NaCl, 0.1 mM MgCl_2_, 0.1 mM CaCl_2_, 5 mM HEPES, titrated to pH 7.3 with NaOH, to which we added drugs, agonists, redox agents, zinc and ion chelators. CTZ stock solution was prepared in DMSO and added at 100 *μ*M to the external solution. Drugs were obtained from Tocris Bioscience, Ascent Scientific or Sigma Aldrich.

### Trapping protocols and chemical modification

To measure the state dependence of trapping in desensitized or resting state in the presence of Cu:Phen, we determined the baseline for activation by 10 mM glutamate in the presence of 5 mM DTT, followed by application of Cu:Phen (10 *μ*M; prepared as described in (Plested and Mayer, 2009)) and the chosen condition in desensitized state in presence of 100 μM Glu in resting without agonist and in the presence of 100 μM CTZ) via the third barrel of the perfusion tool, for different time applications. For all trapping experiments, we quantified the effect of trapping by determining the fraction activated by 10 mM glutamate in 5 mM DTT immediately after trapping as previously described (Lau et al., 2013). The peak current responses after application of Cu:Phen were fit with a single exponential. By back-extrapolating to the end of the Cu:Phen application, we were able to estimate the proportion of receptors that were trapped (Plested and Mayer, 2009). The amplitude of the fit function was the trapped fraction of receptors, and we subtracted this fraction from 1 to get the active fraction (*AF*). For metal bridging experiments, Zn^2+^ was added (10 μM) to the external solution. To achieve zinc-free conditions, we added EDTA to the external solution (10 *μ*M), a potent Zn^2+^ chelator (*K*_D Zn_^2+^ = 10^−16.4^ M). We used the same analysis to determine the active fraction. We applied drugs to outside patches via perfusion tools made from custom manufactured glass tubing with 4 parallel barrels (Vitrocom, Mountain Lakes, NJ) as described in (Lau et al., 2013). The glass was pulled with a finial with of 200 μm, the tip of the tool was etched in hydrofluoric acid and mounted in a piezo electric lever, controlled via a 100 V amplifier. The command voltage was filtered at 100 Hz to reduce vibration of the tool. When we measured the junction potential, the typical 10-90% rise time was 300 μs. For the fast oxidizing experiments, we determined the time that the patch spent in the Cu:Phen condition (third barrel) by measuring open tip currents, calibrating a voltage ramp protocol with different slopes in order to vary the dwell time in the third barrel from 5 to 30 ms (Figure 7A). We measured concentration-response curves for wild-type and the mutant A665W. We obtained the *EC*_50_ from fits to the Hill equation:

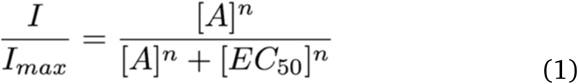

where *n* is the Hill coefficient, *I*_max_ is the maximum response and [*A*] stands for the agonist concentration. Recovery from desensitization for the A665W mutant was measured with a 400 ms conditioning pulse.

**Figure 2.**
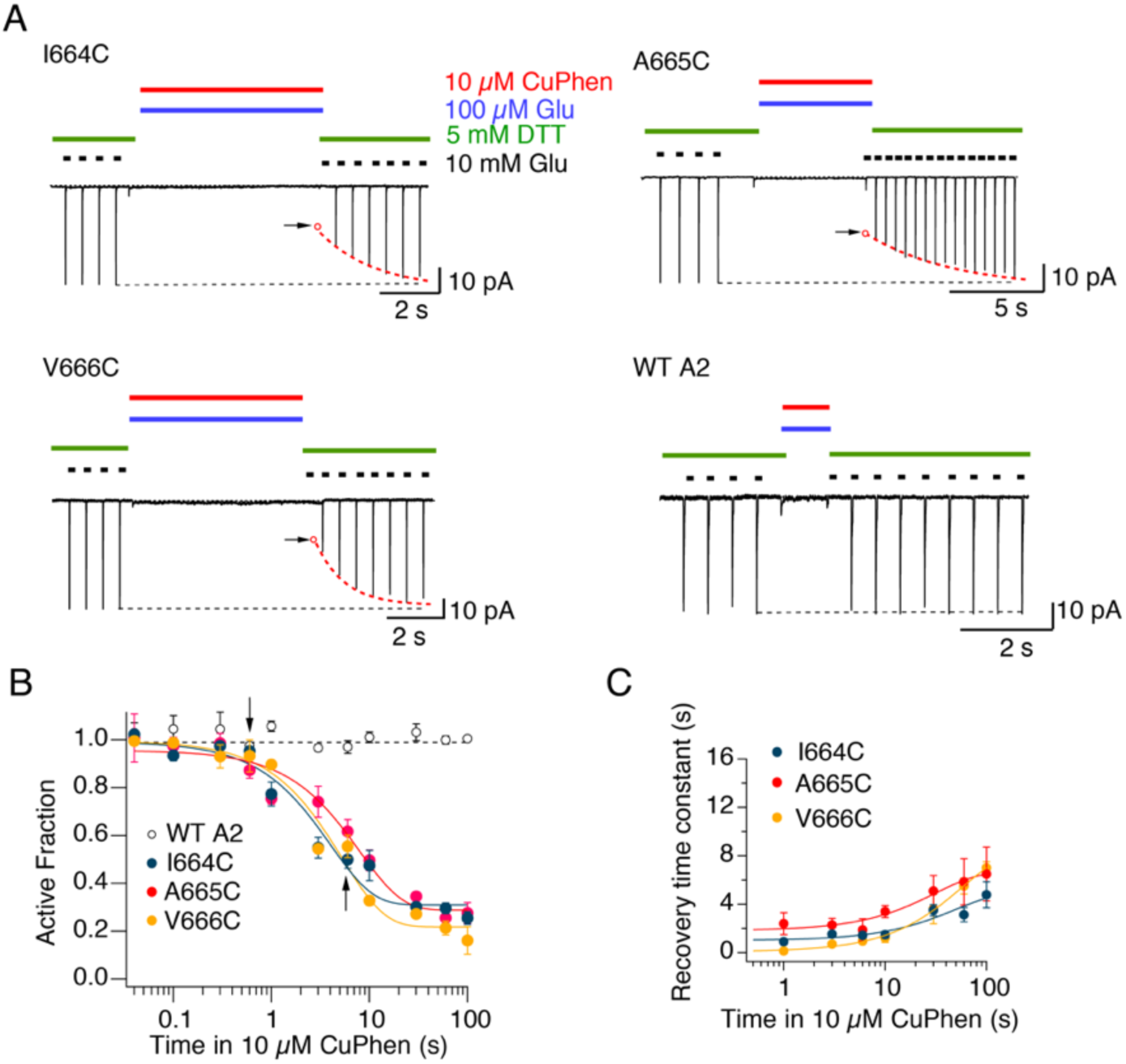
Contact between subunits A and C during desensitization. (A) Typical patch clamp experiments for cysteine mutations at positions 664-666 and WT GluA2. Dotted red line is the exponential fit to recovery of the current in 10 mM glutamate and DTT, following trapping in presence of 10 μM Cu:Phen and 100 μM Glutamate. Open circles with arrow indicate the presumptive back-extrapolated response immediately after trapping. (B) The active fraction of receptors after oxidization in the desensitized state plotted against the trapping interval (continuous lines are exponential fits). The probabilities of no difference in the active fraction after 10 s trapping in 10 *μ*M Cu:Phen were 0.001, 0.0005 and 0.00003 for I664C, A665C and V666C, respectively (compared to WT A2, Student’s *t*-test, *n* = 3-4 patches per point). The arrows indicate the relevant intervals for the traces in panel A. (C) The time constant of recovery after trapping is plotted against the trapping interval for I664C, A665C and V666C, showing a positive correlation (continuous lines are exponential fits; *n* = 3 - 4 patches per point).

**Figure 3.**
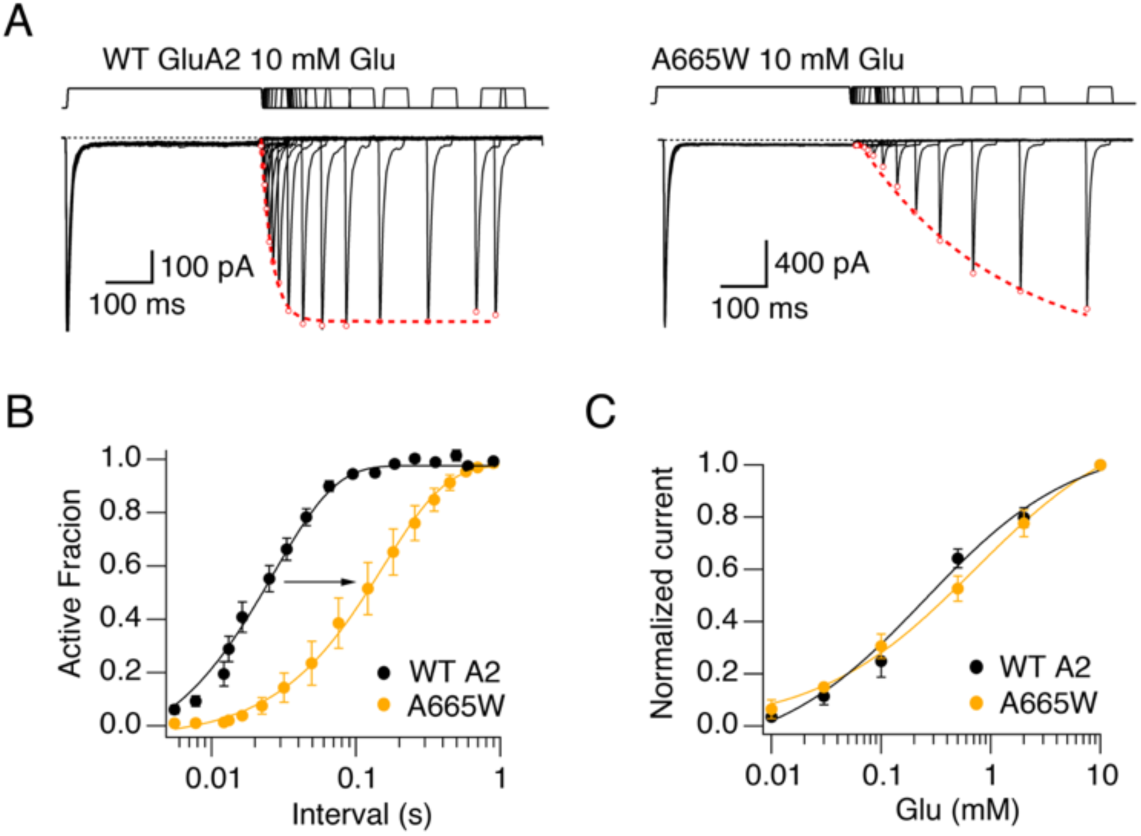
The A665W mutation slows down recovery from desensitization. (A) Currents evoked by 10 mM glutamate from WT GluA2 (left panel) and the mutant A665W (right panel). The two-pulse protocol had a conditioning pulse of 400 ms, followed by a second pulse at increasing intervals (responses are overlaid). Peak currents (red open circles) were fit with Hodgkin-Huxley functions (dashed red lines) (B) Mean of recovery from desensitization for GluA2 WT (black) and A665W (orange). For each interval, the peak of the second pulse is plotted as the active fraction (relative to the first peak) and fit with a Hodgkin-Huxley equation (with slope fixed to 2, see methods). The time constants of recovery from desensitization were 20 ± 2 ms and 155 ± 5 ms for GluA2 WT and A665C respectively (probability of no difference = 0.01, Student’s *t*-test; *n* = 4). (C) Concentration-response curves for WT GluA2 (blue circles; EC_50_ = 330 ± 100 *μ*M), GluA2 A665W (yellow circles; *EC*_50_ = 510 ± 130 μM). The probability of no difference between the *EC*_50_ values was 0.19) (*n* = 3 cells).

**Figure 4.**
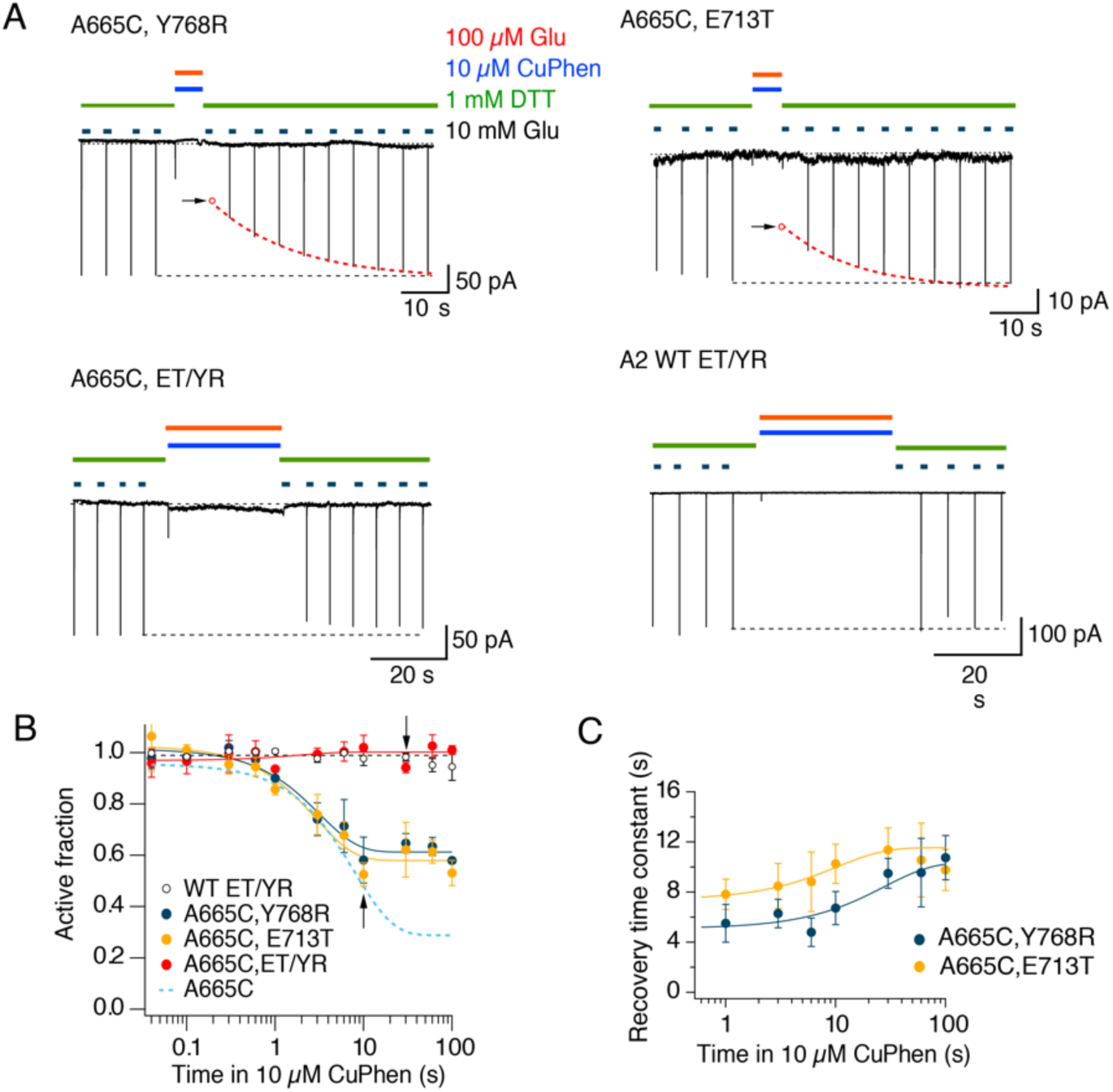
Protection of A665C in deep desensitized states. (A) Typical records showing trapping and recovery for the mutants A665C Y768R (upper left) and A665C E713T (upper right). Arrows indicate the reduction of the current after trapping. The presumptive response (open red circle) was extrapolated from the double exponential fits to the recovery (dashed red lines). The mutant A665C on the ET/YR background (lower left) and the GluA2 ET/YR background (lower right) show no modification. (B) The active fraction of receptors after oxidization in desensitized state (continuous lines are exponential fits) plotted against the trapping interval. The A665C trapping profile (light blue dashed line) and the fit to the ET/YR background (black dashed line) are indicated. The probability of no difference between the active fraction after 10 seconds of application of oxidizing conditions was 0.012 and 0.0008 for A665C, Y768R and A665C, E713T (vs A665C, ET/YR, Student’s t-test, *n* = 3 - 4 patches per point) Arrows indicate intervals for the traces in (A). (C) Time constants of recovery after trapping plotted against the trapping interval for A665C, Y768R and A665C, E713T (continuous lines are exponential fits, *n* = 3 - 4 patches per point).

**Figure 5.**
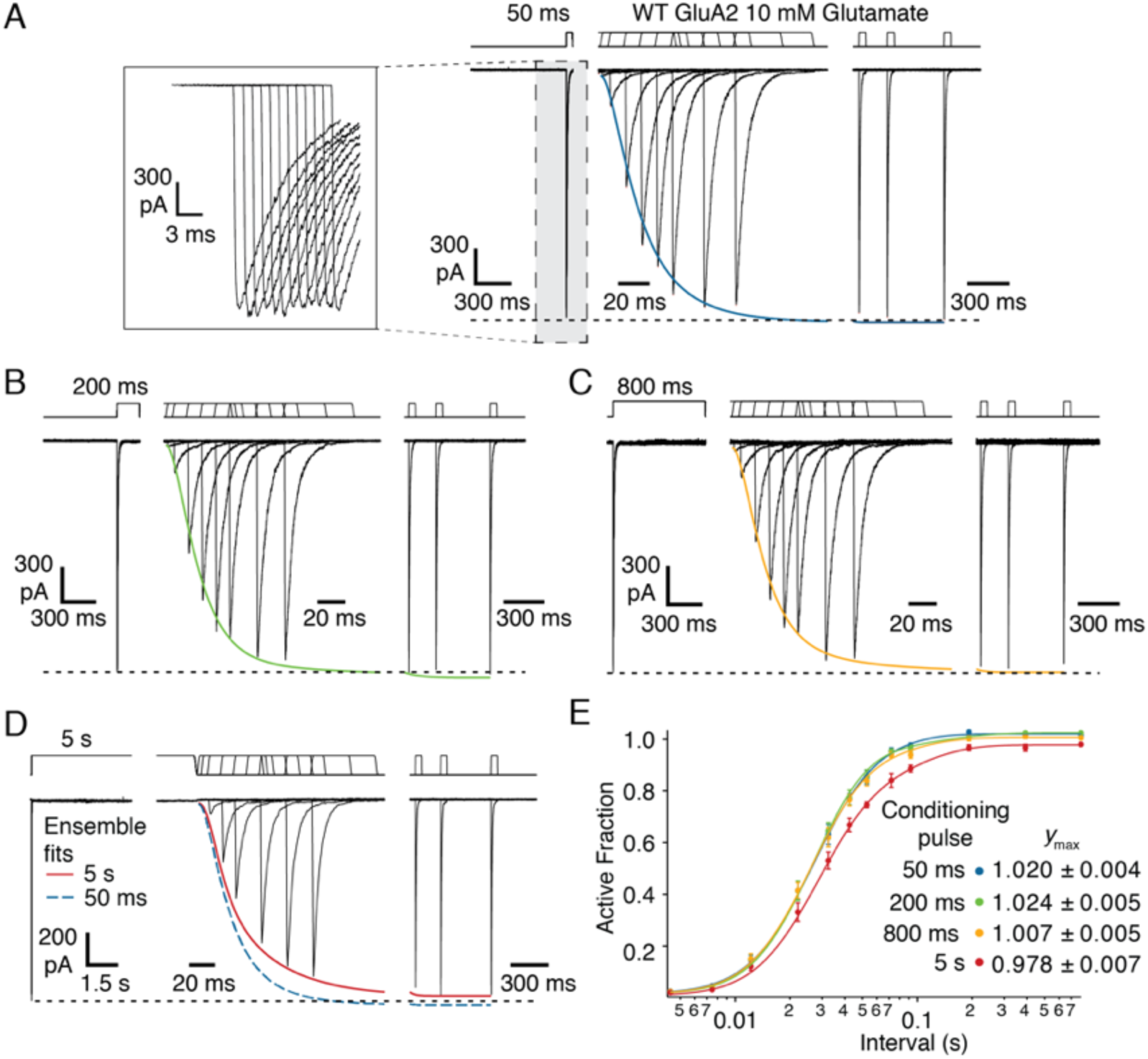
Recovery of wild-type GluA2. (A -D) Two pulse protocols with conditioning pulses of 50 ms to 5 s. Inset (from grey boxed area) shows the typical consistency of responses to the conditioning pulse; responses were also highly consistent in their timing but are drawn displaced here. Ensemble fits to data with the two component H.-H. function (equation 3, see methods; conditioning pulses of 50 ms blue, 200ms green, 800 ms orange, 5 s red) are overlaid on traces from immediate and late recovery responses (note changes of abscissa scale). Dashed black line indicates the responses to the conditioning pulse. In panel D, the 50 ms ensemble fit is included as a dashed blue line. Note that the 5 s ensemble fit (red) did not reach the amplitude of the response to the conditioning pulse. The interval between episodes was typically 1 s. (E) Ensemble fits with equation 3 (H.-H. with 2 components). The slope of the fast component was fixed to 4 and the slope of the slower component was fixed to 2. Fitted maxima are shown. For the 5 s conditioning pulse, *a*_1_ = 0.7, *k*_1_ = 71 s^−1^, *a*_2_ =0.27. *k*_2_ = 19 s^−1^ with maximum 0.978 as shown. Alternative fits and fit parameters with errors are shown in Supplementary Figures 1 & 2 and Supplementary Tables 1 & 2.

**Figure 6.**
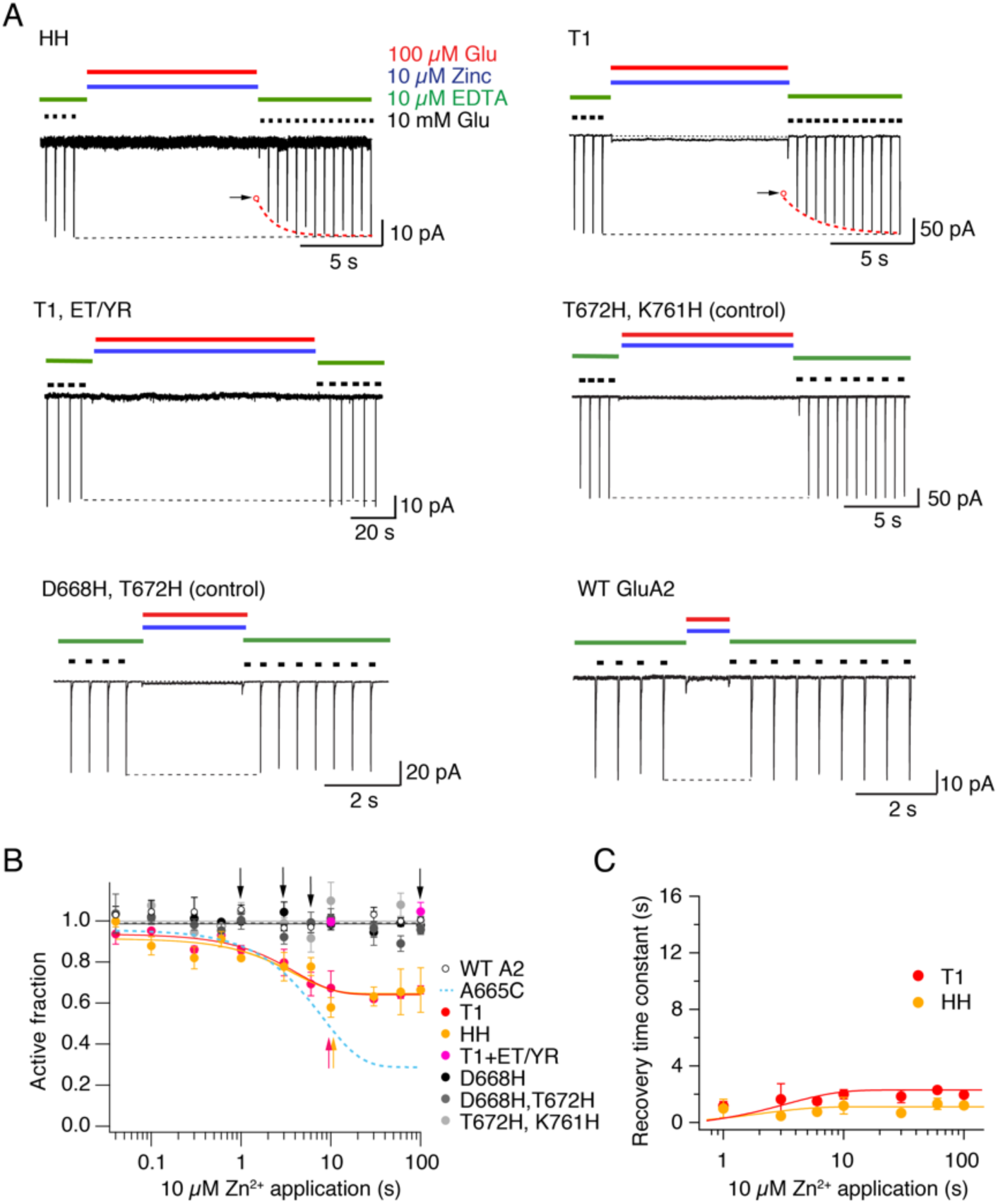
Lateral movements of the LBDs during desensitization. (A) Typical records for trapping and recovery from zinc trapping for the mutants HH (upper left panel) and T1 (upper right). Arrows indicate reduction of the current after trapping; dotted lines are double exponential fits to the recovery after trapping. The mutant T1 ET/YR shows no inhibition after 100 s exposure to zinc. Controls, such as the double mutants D668H, T672H and T672H, K761H also showed no modification, behaving like WT GluA2. (B) The active fraction of receptors after Zn^2+^ exposure in desensitized state (continuous lines are exponential fits) plotted against the interval. The probability of no difference (Student’s *t*-test) to the WT A2 (open circles) after 10 seconds of Zn^2+^ was 0.018 for T1 (red circles) and 0.001 for HH (yellow circles). Controls and T1 ET/YR were indistinguishable from WT A2. Arrows indicate intervals for the traces in (A). (*n* = 3 - 4 patches per point). (C) The time constants of recovery after Zn^2+^ trapping plotted against the trapping interval for T1 and HH (*n* = 3 - 4 patches per point).

**Figure 7.**
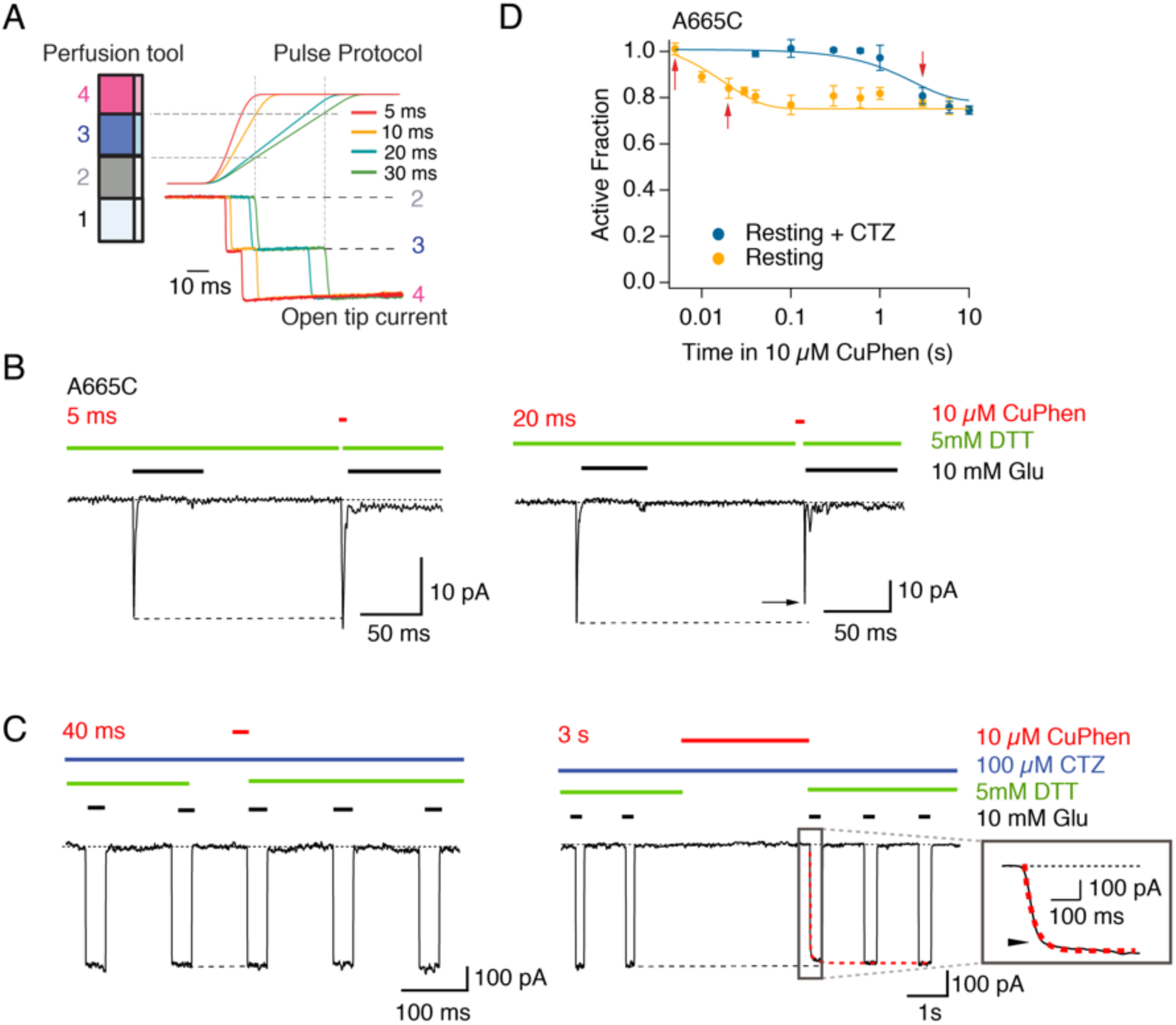
Rapid resting state desensitization. (A) Using the four-barrel fast perfusion system, and switching from barrel 2 to 4, we applied oxidizing conditions for intervals as brief as 5 ms, with a voltage ramp command to the piezo lever (right panel). The open tip junction current shows that the pipette spends between 5 and 30 ms in the 3^rd^ barrel outflow (blue). Dashed lines indicate switch times for the 20 ms exposure. (B) Patch clamp experiments showing Cu:Phen (10 *μ*M) exposures of 5 ms (left panel) and 20 ms (right panel) for the mutant A665C in the resting state. Arrows indicate reduction of the current after modification. (C) Patch clamp experiments showing applications of Cu:Phen 10 *μ*M for 40 ms (left panel) and 3 s (right panel) to. the mutant A665C in resting state with CTZ (100 *μ*M). The inset shows the current profile during recovery: the arrow indicates reduction of the current after trapping. (D) The active fraction of receptors after oxidization in desensitized state (continuous lines, exponential fit) is plotted against the trapping interval. Arrows indicate intervals for the traces in (B and C). The probability of no difference between the active fraction after trapping in resting state without and with CTZ for A665C, after 10 seconds of application of oxidizing conditions was 0.003 (*n* = 3 - 6 patches per point).

### Recovery from desensitization

To examine the recovery from desensitization for wild-type GluA2, patches containing hundreds of receptors were conditioned with applications of 10 mM glutamate for 50, 200 and 800 ms and 5 seconds. A test glutamate pulse was delivered at 12 different time points between 2 and 790 ms after the conditioning pulse. The protocols with different durations of the first pulse were randomly initiated for each patch and repeated, if the patch was stable enough.

The timing and amplitude of peak currents and rise times of all peaks were measured in Axograph. Recordings from 23 patches from different cells were used for further analysis, except for the 5-second conditioning pulses. Since the duration of these measurements was long and the run-down of the current was often substantial, only measurements from 4 patches could be completed before the patch was lost, and had sufficiently good quality for the whole set of records, namely 10-90% rise times of the glutamate response < 500 μs, and a stable baseline with fluctuations less than 10% of the peak current.

The response following a long (5 s) conditioning pulse was corrected for the slow rundown of current amplitudes caused by either accumulation of receptors into electrically-isolated parts of the patch (Suchyna et al., 2009), or accumulation of receptors into non-functional states. A linear function was fitted to currents and times of the response to the conditioning (1^st^) pulse from each episode, to extrapolate the expected maximum response at the time of each test (2^nd^) pulse. For each interval, the normalized responses were averaged and the standard deviation was used for fitting in Igor Pro (Wavemetrics).

We and others have previously fitted recovery from desensitization with a Hodgkin-Huxley type recovery curve (Hodgkin and Huxley, 1952):

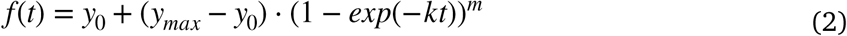

where *k* is the rate of recovery, *m* is the slope, *y*_max_ and *y*_0_ are the maximum and minimum, respectively, and *t* is the interval between pulses. We also did this here for the data in Figure 3A. However, for the recovery following conditioning pulses of longer durations (Figure 5), this function was insufficient because it could not describe the intermediate phases of recovery. These data could only be well fit by a function that was the sum of two Hodgkin-Huxley terms (with rates *k*_1_ and *k*_2_, slopes *m*_1_ and *m*_2_):

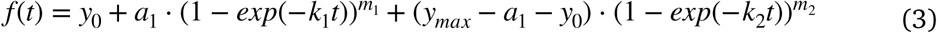

To establish how unique the description by this two component H-H function was, we tried several other different fit functions (3 Component H.-H. function, Sigmoid, Hill) and varied the fit parameters (see Supplementary Tables 1 & 2 and Supplementary Figures 1 & 2). Time constants where quoted are reciprocals of rate constants.

### Statistical Analysis

All *p*-values were determined by a two-sample unpaired Student’s *t*-test. The spread of the data where indicated is the standard deviation of the mean.

## Results

### An inter-dimer interface forms in desensitized state

To place intersubunit bridges in the context of the conformational changes that drive desensitization, we first compared the LBD layers of the apo state structure of GluA2 (PDB code: 4U2P; Dürr et al., 2014) to three candidate desensitized state structures. We modelled the A665C mutation into these structures and measured the putative distances between the Sulphur atoms of the cysteine. In the resting ligand-free structure the SG-SG distance is 9 Å, partly due to the open angle adopted by the dimers but mainly as a consequence of the distance of the A and C subunits from the central axis of the receptor and ion pore. The structure of the GluA2 receptor in complex with the high-affinity full agonist L-quisqualate and the accessory subunit GSG1L (GluA2-2xGSG1LQuis. PDB: 5VHZ; Twomey et al., 2017) presents LBD dimers with broken local 2-fold symmetry, with disrupted interfaces between the dimers due to rotation of 31° and translation of 6 Å of the upper D1 lobes. Despite these motions, the closest approach of modelled SG-SG distance remains 9 Å (Figure 1). In The GluA2-TARP γ-2 Quis (PDB: 5VOV; Chen et al., 2017). shows similar rotations and translations within each of the LBD dimers. Placing the A665C mutations in subunits A and C of this structure allows the SG groups to be only 5 Å distant, within striking distance of forming a disulfide bridge suggesting the TARP-bound desensitized state is quite compact; this concept was discussed at length in (Baranovic and Plested, 2018). A yet more dramatic change can be observed in the homologous GluK2 structure in the desensitized state (2S,4R-4-methylglutamate PDB: 5KUF; Meyerson et al., 2016) which shows a rotation of the subunits D and B of 125°, producing pseudo four-fold symmetric arrangement of the LBDs. Similar large-scale disruptions of the LBD layer were observed in single particle analyses of GluA2 without auxiliary subunits (Dürr et al., 2014; Meyerson et al., 2014; Meyerson et al., 2016). As a consequence of this movement the residue T670 (equivalent in GluK2 to A665 in GluA2) is buried between the new interfaces formed between adjacent subunits (Figure 1). For the same four candidate structures, we measured the distances between the residues that form the site of the T1 lateral Zn^2+^ bridge (Baranovic et al., 2016) built from introduced histidines (D668H T672H K761H; Figure 1C). The intersubunit distances are too great to predict zinc bridging for any of the candidate structures. The closest approach was for GluA2-TARP γ-2 Quis (5VOV), where the CA-CA distances were 12-13 Å (for K761 to D668 or T672). Overall, no candidate desensitized arrangement would support a Zn^2+^ bridge if histidine residues were placed at these positions.

We reasoned that if we attempted to crosslink the diagonal A-C interface during desensitization, we could determine at which point in the desensitization reaction (either early or late) that these two subunits come together. Likewise, we expected that the lateral interface should not be readily accessible to desensitized receptors, unless a spectrum of different desensitized states are sampled. We used well-characterized cysteine substitutions at three positions in the FG-loop (I664, A665 and V666) (Lau et al., 2013; Salazar et al., 2017; Baranovic and Plested, 2018). Each mutant was tested for its crosslinking potential in 100 μM glutamate. This concentration was based on the concentration dependence of desensitization of the WT receptor, which reaches full desensitization at 100 μM (Plested and Mayer, 2009), with minimal activation. We used three barrels of a quadruple barrel fast perfusion system that enabled application of 100 μM glutamate in the presence of 10 μM Cu:Phen with < 10 ms resolution. We observed a dramatic reduction of the current after exposure to oxidizing conditions for the mutants I664C, A665C and V666C (Figure 2 A). The untrapped, active fraction (*AF*) was measured for different intervals of exposure to oxidizing conditions in desensitized state. After 10 seconds of oxidizing conditions the reduction of the AF reached a plateau for I664C, A665C and V666C (47 ± 6 %, 49 ± 4 % and 32 ± 2 %, respectively (Figure 2B). The kinetics of recovery after trapping indicate how stable this interface is in the desensitized state. The time constants of recovery after trapping for I664C, A665C and V666C were 1.4 ± 0.2 s, 3.3 ± 0.6 s and 1.2 ± 0.4 s respectively after 10 seconds in Cu:Phen (Figure 2 C). Strikingly, with the longer exposure to Cu:Phen, the three mutants I664C, A665C and V666C each showed increased stabilization of the trapped state. We fitted the increase in the recovery time constant against exposure time with a single exponential to obtain the asymptotic maximum recovery time constants of 43 ± 13 s, 38 ± 19 s and 43 ± 5 s for I664C, A665C and V666C, respectively (Figure 2 C). These results indicate that the A-C interface can be trapped in at least two desensitized states, one of which recovers rapidly. However, with long exposures, there is a progressive adoption of a state or set of trapped states that are considerably more stable.

### A point mutant at the A-C interface slows recovery from desensitization

According to previous reports, the introduction of a cysteine in position A665 alters recovery from desensitization in a redox sensitive manner (Yelshanskaya et al., 2016). We hypothesised that the introduction of a bulky residue like tryptophan in position A665 should alter recovery, if this interface forms during the desensitization process. To test this, we conditioned patches with an application of glutamate (10 mM for 100 ms), and then a second application was delivered at varying times after the conditioning pulse. In these experiments, WT GluA2 recovered from desensitization with a time constant of 20 ms, as previously reported (Carbone and Plested, 2012) In contrast, we observed a dramatic delay of desensitization recovery of more than 7-fold for the mutant A665W (τ_rec_ = 155 ± 5 ms; Figure 3 A). We constructed a dose response curve for glutamate and observed little difference in the apparent affinity for glutamate (A665W *EC*_50_ = 510 ± 130 μM compared to WT *EC*_50_ = 330 ± 100 μM, with *p* of no difference 0.19) (Figure 3 B). We therefore ruled out the possibility that the change in recovery was due to an increase in affinity for glutamate in the A665W mutant. This observation further supports the idea that this inter-subunit interface forms during the entry to or exit from desensitization.

### Adoption of deep desensitized states protect against crosslinking

We reasoned that if the stable disulfide trapping we detected were unique to the desensitized state, then promoting desensitization should promote trapping and/or slow recovery. However, additional desensitized states might exist that are not readily disulfide linked by cysteines at the A-C interface.

Such states would perhaps resemble the four-fold symmetry of the GluK2 structure, or more generally, would stably hinder the approach of cysteines at the otherwise proximal A-C interface because of a substantial conformational change. We tested the formation of inter-subunit crosslinks in a putative deep desensitized state using a mutants that strongly stabilizes the desensitized state (E713T, Y768R), with a recovery time constant of 1.1 ± 0.2 s (Carbone and Plested, 2012). When we introduced a cysteine in position A665 in the individual backgrounds E713T and T768R, we observed less profound trapping than for the A665C mutant alone, with a reduction of the active fraction of 46% for A665C, E713T and 42% for A665C, Y768R after 100 s of application of Cu:Phen (Figure 4 A and B). Both mutants showed slower recovery after trapping for A665C, E713T of 10.7 ± 1.7 s and 9.7 ± 1.6 s for A665C, Y768R (Figure 4 C). Again, longer exposures to Cu:Phen drove adoption of a yet more stable trapped arrangement. Fitting a single exponential to the recovery time constants vs the time in oxidizing conditions, we determined the asymptotic limiting time constants of recovery of 27 ± 10 s and 6 ± 1 s, (for A665C, E713T and A665C, Y768R, respectively) (Figure 4 C). In dramatic contrast to these results, the triple mutant A665C ET/YR showed absolutely complete protection from trapping, exhibiting a similar profile to the WT GluA2 ET/YR mutant (Figure 4 A). This observation suggests that this GluA2 mutant, which exhibits very stable desensitization, can adopt yet another deep desensitized state in which the interface between subunits A and C is absent. Such conformation is consistent with either the GluA2-2xGSG1LQuis structure, or the GluK2 Cryo-EM structure where the equivalent residue for A665C is buried in intersubunit interfaces (Figure 1).

### Long exposures to glutamate promote entry to stable desensitized states

From these results, we predicted that the progressively greater stability of trapped receptors following long exposures to desensitized conditions should derive from the selective adoption of more stable desensitized states. However, the wild-type homomeric GluA2 receptor is known for its rapid and complete recovery from desensitization (see for example Figure 3A). To resolve this paradox, we tested if the rate of recovery from desensitization was sensitive to the duration of the conditioning pulse in two-pulse recovery experiments. In each record, a patch containing hundreds of wild-type GluA2 receptors was first conditioned with an application of glutamate (10 mM) for 50, 200 and 800 milliseconds and 5 seconds. A second glutamate pulse was delivered at varying times after the conditioning pulse. (2 - 790 ms; Figure 5). For each patch, we made series of recordings with the same conditioning pulse, but different intervals, and then repeated the protocol with a different conditioning pulse length. Following short conditioning pulses, we observed prototypical fast recovery from desensitization for GluA2. The recovery following 50 ms and 200 ms conditioning pulses could be quite well described with a single H.-H.-type function with a time constant of 20 ms (with slope between 2.5 and 3; Supplementary Figure 1B). However, in the same patch, a 5 second conditioning pulse of glutamate slowed the recovery profile in 2 distinct ways (Figure 5E). Firstly, the fastest component, with a time constant of about 14 ms, was only ∼70% of the total amplitude (Supplementary Table 1). Second, a small (∼5%) but very slow (∼1 s) component meant that recovery at the end of our protocol (∼700 ms interval) was always incomplete. Comparison of the recovery after 800 ms and 5 s conditioning pulses revealed both had a small intermediate component (∼20%) with time constant ∼50 ms (Supplementary Table 1).

We also observed a slow component of recovery when using 10 s or 30 s conditioning pulses, but it was hard to quantify because the recovery protocols using such long conditioning pulses necessarily lasted for 5 – 10 minutes, over which time even stable patches ran down and gave spurious responses. Taken together, these observations emphasise that wild-type GluA2 receptors can adopt a range of desensitized states with different stabilities, including very stable desensitized states.

### Lateral shifts occur during desensitization

The GluA2-TARP γ2 Quis (5VOV) structure (Chen et al., 2017) shows a compact packing of the lateral interface of the subunits A&B and C&D of the LBDs whereas in the GluA2-2xGSG1LQuis (5VHZ) structure (Twomey et al., 2017) this interface is clearly absent (Figure 1C). In order to analyse if this if this interface occurs in early-desensitized states that can be adopted over millisecond time scales, we used a metal ion trapping approach. Previously, we engineered a pair of histidine mutants T1 (D668H, T672H, K761H) and HH (D668H, K765H), which can coordinate Zn^2+^ between subunits A & B or C & D at intermediate and high concentrations of glutamate, with cyclothiazide present to block desensitization (Baranovic et al., 2016). Using these mutants, we detected the formation of the lateral interfaces in desensitized state applying 100 μM glutamate in presence of 10 μM Zn^2+^. The mutants T1 and HH showed a reduction of the active fraction after 1 second of the application of Zn^2+^ in desensitized state, with a reduction of the AF of 33% for T1 and 23% for HH plateauing after 100 s of Zn^2+^ application (Figure 5 A). With time constants of recovery after trapping for T1 and HH of 1 ± 0.08 s, and 0.6 ± 0.08 s respectively after 10 seconds in zinc. The asymptotic time constants of recovery in the limit of long Zn^2+^ exposures were 4 ± 0.6 s and 2 ± 0.8 s, (for T1 and HH, respectively; Figure. 5 A).

We hypothesised that the conformation trapped by the T1 lateral bridge is absent in the deep desensitized states promoted by the ET/YR mutation. To test this hypothesis, we inserted the triple histidine mutant T1 (D668H, T672H, K761H) in the mutant (E713T, Y768R) background and tested its sensitivity to zinc in desensitized states. As for the A665C mutant, we did not detect the formation of T1 lateral interfaces in the presence of the ET/YR mutation. The T1 ET/YR mutant was as insensitive to 10 and 100 s application of 100 μM glutamate in presence of 10 μM Zn^2+^ as WT GluA2 (Figure 5 A). Double and single histidine mutant controls were also unaffected by Zn^2+^ application (Figure 5 A), confirming the specificity of the trapping of the mutants T1 and HH. These experiments show that lateral shifts occur during desensitization, and the fast formation of bridges over tens of milliseconds suggests that these shifts occur during the 1^st^ steps of the desensitization pathway. Additionally, we observed no lateral interface formation in the deep desensitized state.

### Resting state desensitization is rapid and reversible

The differences between the NW-bound, putative desensitized structure (4u4f) (Yelshanskaya et al., 2014), the apo structure(Dürr et al., 2014) and the MPQX-bound state (Twomey et al., 2016; Zhao et al., 2016) are subtle, with little to no change in the distance between subunits A and C at A665C between (7 to 9 Å). All these structures have preserved intra-dimer active D1-D1 interfaces, and previous work showed that unbound and singly bound receptors can undergo desensitized transitions (Robert and Howe, 2003; Plested and Mayer, 2009). This raises the question as to whether resting receptors can be trapped at the lateral interface as they desensitize. To investigate this point, we studied the formation of an inter-dimer crosslink between the residues A665C in the absence of ligand. Initial experiments suggested crosslinking that was effectively instantaneous, using our regular protocols. To make the briefest application of Cu:Phen possible, we programmed a ramp stimulation for the perfusion tool, as illustrated in Figure 6A. With this protocol, we could apply oxidizing conditions for less than 5 ms. Exposing the mutant A665C to oxidizing conditions (10 μM Cu:Phen) in absence of any agonist, we observed trapping of the A665C mutant that developed over about 10 ms, reducing the fraction of active receptors by about 20% (Figure 6B). This trapping was comfortably the fastest that we have observed in the AMPA receptor (Lau et al., 2013; Baranovic et al., 2016; Salazar et al., 2017). To investigate whether this trapping requires breaking of the active dimer interface, we constrained the interface by exposing patches to 100 μM cyclothiazide (CTZ). There was an approximately 1000-fold delay in the formation of the diagonal disulfide between subunits A and C in resting state in the presence of CTZ, with a reduction of the AF of 20% only after 100 s application of Cu:Phen (Figure 6 C and D). The time constant of recovery after trapping in resting conditions was 380 ± 150 ms, but for resting + CTZ the recovery was much faster, 30 ± 5 ms (after 10 seconds in Cu:Phen). These results show that the receptor transits between active (D1 intact) and desensitized-like (D1 broken) states on a more rapid timescale at rest than previously thought (Plested and Mayer, 2009).

## Discussion

The idea that the interaction of the neurotransmitters with receptors encompasses more than a simple binding interaction followed by activation was a major step in receptor theory (Katz and Thesleff, 1957). It is now known that receptors in the brain have multiple active and inactive “desensitized” states. For example, the acetylcholine receptor presents at least four desensitized states (Elenes and Auerbach, 2002), while for the BK potassium channel, which has multiple Ca^2+^-binding sites, desensitized configurations could represent up to 120 different states (Magleby, 2003). Previous work has emphasised that native AMPA receptors, likely in complex with auxiliary subunits, have multiple desensitized states (Patneau and Mayer, 1991; Colquhoun et al., 1992) as do TARP-associated receptors expressed recombinantly (Levchenko-Lambert et al., 2011). Multiple components in the recovery of AMPA receptors expressed in cell lines are detectable but less pronounced (Bowie and Lange, 2002; Robert and Howe, 2003). However, the GluA2 (Q) homomer that we worked on here, and for which the majority of structural studies were completed, was until now widely reported to have a rapid and monotonic recovery from desensitization (see Figure 3) (Zhang et al., 2006; Weston et al., 2006; Carbone and Plested, 2012). We mapped inactive conformations of GluA2 over timescales from milliseconds to minutes with metal bridges and disulfide bonds that trap transient intersubunit interactions. This approach facilitated the detection of 3 distinct classes of desensitized states with glutamate, and a fast inactive state at rest (Figure 8A).

**Figure 8.**
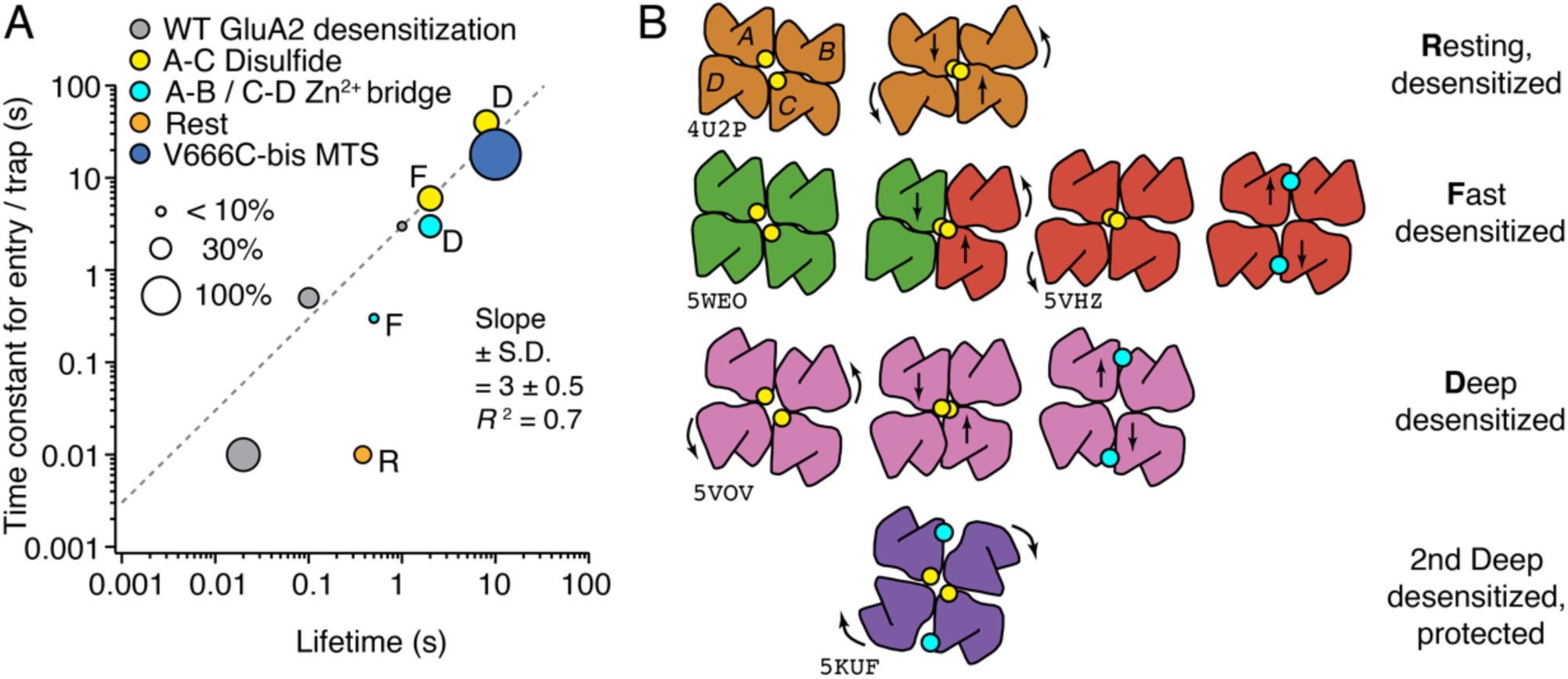
Models of trapping in desensitized states. (A) The time constants of trapping and entry to desensitization are plotted against desensitized or trapped state lifetimes. The symbol sizes represent the fraction of receptors trapped as indicated. Entry time constants depend on condition (zinc concentration, bis-MTS concentration), hence the linear relation has little physical meaning. Entry time constants for trapping were estimated from midpoints of exponential fits to lifetimes. Lettering indicates presumptive ‘R’esting, ‘F’ast and ‘D’eep classes from panel B. The V666C-bis MTS point (dark blue) is from 1 *μ*M bis-MTS trapping (Baranovic and Plested, 2018). (B) Putative LBD arrangements in 4 distinct classes of desensitized states identified by crosslinking (plan view). Subunits A-D are arranged as marked on resting state. Published structures are indicated by PDB codes. Diagonal disulfide bonds are indicated by paired yellow circles, and the lateral Zn^2+^ bridge by a cyan circle (as in A). Fast desensitization can probably occur with one dimer active (green) and one dimer desensitized (red), after only small movements. Multiple components of trapping for both disulfides and zinc bridges require multiple states, here represented by the “Fast” and “Deep” desensitized rows (lettering from panel A indicated in bold here). In the 2^nd^ deep desensitized state, neither the A-C disulfide nor the lateral bridge can form.

We observed a positive correlation between the time of application of oxidizing conditions and the time constant of recovery after trapping for disulfide bonds at the A-C subunit interface, and lateral zinc bridges. The simplest explanation is that prolonged exposure drives entry to at least two desensitized states, and the least frequently accessed are the most stable (see Figure 8A). The recovery from trapping in these desensitized states was dramatically slower than for the same mutants trapped in active states (Baranovic et al., 2016; Lau et al., 2013). Most surprisingly, by favouring slow recovery by introducing mutations in the D2 lobe of the LBDs, we could access a further, conformationally distinct, stable desensitized state that was immune to crosslinking at the A-C interface and that may or may not be physiological. It is tempting to consider these three distinct states as progressively profound conformational changes in the LBD layer, with more stable desensitized states corresponding to those seen in some structural biology experiments. In these experiments, ligand exposures are for technical reasons minutes or hours. In Figure 8B, we outline a scheme to link the states identified from their crosslinking behaviour to possible LBD arrangements.

Although crosslinks do not define geometry uniquely, we note that they do report a minimum level of complexity in the conformational and dynamic space of GluA2. The three time constants we identify for wild-type GluA2 (in glutamate) may correspond to the “fast”, “deep” and “protected” classes (Figure 8B). Assuming this relation would imply that disulfides and zinc bridges trap these states about 100 times slower than the native states are entered (at the concentrations of Cu:Phen and Zn that we used), and extend the lifetime of the trapped states by about 100-fold. A back of the envelope calculation applying this logic to the observed time constants for the resting state trapping (entry ∼10 ms, lifetime ∼ 400 ms) would give the true resting state D1 dimer desensitization with time constant for entry of about 100 *μ*s, and a lifetime of ∼4 ms. Providing some support for these estimates is the separate observation that trapping with bifunctional MTS reagents can be accelerated 50-fold by simply increasing the reagent concentration (Baranovic and Plested, 2018). In this and other studies, we typically trap receptors in gentle (read, slow) conditions to reduce non-specific crosslinking

Even though prolonged oxidation can lead to promote trapping by disulfides in stable inactive states, key weaknesses of this line of approach are that the presence of the bridges themselves could drive non-physiological conformations, and the trapping bridges necessarily contribute to the lifetime of the trapped states. To address these points, we exposed wild-type GluA2 receptors to long applications of glutamate in two-pulse protocols, and could detect slow components of recovery when we used conditioning pulses of 800 ms or longer. About one third of the population recovered either with an intermediate recovery rate about 4 times slower than the majority, or on a timescale longer than the pulse protocol (>1 s). With the brief conditioning pulses that we and other investigators have routinely used (Figure 3 and (Bowie and Lange, 2002; Zhang et al., 2006; Carbone and Plested, 2012), these slow components are either very small or absent. The Hodgkin-Huxley type functions that describe recovery of GluA2 well are usually fixed to have a slope of 2. Intriguingly, a good description of the early phase of recovery required slope exponents > 2 (see Figure 5, Supplementary table 1 and Supplementary Figure 1). Fixing the slope to 4 offered a marginal improvement in the goodness of fit compared to a slope of 3. This observation is consistent with 3 or more independent particles being involved in the first recovery phase (Hodgkin and Huxley, 1952). A more qualitative observation is that, for conditioning pulses of 5 s or longer, we always observed rundown of the response. Although there are multiple explanations of rundown (see Methods), one source could be the irreversible accumulation of receptors into non-functional states. Quantitative measures of receptor conformation during such experiments (for example, from spectroscopy) may be able to provide information in this regard.

The desensitized state structure stabilized by GSG1L does not support formation of the disulphide bond between A665C residues in subunits A and C (Figure 1). This observation reinforces the idea that there are multiple desensitized states with common attributes, but that some aspects of LBD geometry might be unimportant. Desensitized states in the AMPA receptor may correspond to any number of configurations where the braced, active dimer arrangements are absent. Dissociation of a single active dimer is enough to desensitize the receptor (Robert and Howe, 2003). However, the overall configuration of the four LBDs might otherwise be compact. Auxiliary proteins with very different geometries (for example, TARPs and Shisa variants) seem to have distinct effects on the lifetime of the AMPA receptor desensitized state (Priel et al., 2005; Farrow et al., 2015; Eibl and Plested, 2017), and this could be because they stabilize different LBD arrangements in desensitized states

Structures of GluA2 in the apo ligand free (Dürr et al., 2014), and the MPQX-bound state (Twomey et al., 2016; Zhao et al., 2016), do not support a contact between subunits A and C in resting state. Yet inactive states could be trapped by the A665C disulfide bond at “rest” within 10 milliseconds. Therefore, these measurements place an upper limit of the timescale of latent rearrangements of the dimer interface. The S729C mutant forms a precedent for these observations (Plested and Mayer, 2009), but for that mutant, resting state trapping was about 700-fold slower. Using CTZ, to stabilize the D1-D1 interface, we could massively slow trapping in resting state, suggesting that D1 dissociation is rapid and regular (Gonzalez et al., 2010). It is likely that reformation of the interface at rest is at least as fast, otherwise the majority of receptors should simply desensitize upon binding glutamate. A second, less likely, effect of CTZ would be to reduce the conformational dynamics of lower lobe of the LBD (where A665C is located) in the resting state. In presence of partial agonists, twisting motions and other degrees of freedom that have been reported (Fenwick and Oswald, 2008; Ahmed et al., 2011; Birdsey-Benson et al., 2010), and these may be affected by CTZ binding.

Our results are consistent with a previous report that shows differences in the rates of entry and recovery from desensitization in A665C receptors between oxidizing and reducing conditions (Yelshanskaya et al., 2016). The introduction of tryptophan in position A665 produces a slow rate of recovery from desensitization of more than 7-fold. Therefore, a movement involving a close contact between subunits A & C at the loop between helices F and G of the LBDs appears necessary for fast recovery from desensitization.

In summary, our experiments provide insight into the conformations and kinetics of AMPA receptor desensitization. Particularly, this work suggests a hierarchy of AMPA receptor desensitized states. It seems likely that the more dispersed configurations of the extracellular domains detected in some structural biology experiments correspond to slowly-attained states. These states may be relevant in pathologies or for aspects of AMPAR biology outside of fast excitatory synapses. However, compact desensitized arrangements of the LBD layer, probably like those stabilized by auxiliary proteins observed in Cryo-EM experiments, are rapidly attained by AMPA receptors, and on a timescale relevant for desensitization in the brain.

## Acknowledgements

This work was supported by the ERC grant 647895 “GluActive”, the Deutsche Forschungsgemeinschaft (DFG, German Research Foundation) under Germany’s Excellence Strategy – EXC-2049 – 390688087 and the DFG grant PL619.1 (to A.J.R.P.). We thank J. Baranovic for comments on the manuscript. H.S. was the recipient of a Long-Term Fellowship from Human Frontier Science Program (HFSP).

## Author contributions

All authors performed experiments, analysed data and wrote the paper.

