## Supplemental Tables 1 & 2, Supplemental Figures 1 & 2 for "Measurements of the timescale and conformational space of AMPA receptor desensitization"

**Supplementary Table 1 Hodgkin-Huxley type fits to GluA2 recovery from desensitization with 1, 2 or 3 components.**

| Figure | Function | Duration of conditioning pulse | Goodness of fit ($\chi^2$) | Parameters | | | | | | | | | |
|---|---|---|---|---|---|---|---|---|---|---|---|---|---|
| | | | | $y_0$ | $a_1$ | $a_2$ | $k_1$ | $k_2$ | $k_3$ | $m_1$ | $m_2$ | $m_3$ | $y_{max}$ |
| S1A | H.-H. 1 component | 50ms | 44 | 0.000 ± 0.004 | | | 43 ± 1 | | | *2* | | | 1.022 ± 0.003 |
| | | 200ms | 36 | 0.000 ± 0.006 | | | 44 ± 1 | | | *2* | | | 1.022 ± 0.004 |
| | | 800ms | 41 | 0.000 ± 0.006 | | | 44 ± 1 | | | *2* | | | 1.007 ± 0.004 |
| | | 5s | 153 | 0.000 ± 0.002 | | | 34 ± 1 | | | *2* | | | 0.983 ± 0.006 |
| S1B | H.-H. 1 component | 50ms | 13 | 0.010 ± 0.006 | | | 54 ± 3 | | | 2.8 ± 0.3 | | | 1.017 ± 0.003 |
| | | 200ms | 14 | 0.0011 ± 0.009 | | | 52 ± 3 | | | 2.6 ± 0.3 | | | 1.018 ± 0.004 |
| | | 800ms | 18 | 0.00 ± 0.01 | | | 52 ± 3 | | | 2.5 ± 0.3 | | | 1.001 ± 0.004 |
| | | 5s | 29 | 0.002 ± 0.006 | | | 46 ± 2 | | | 2.8 ± 0.2 | | | 0.967 ± 0.006 |
| S1C | H.-H. 2 component | 50ms | 940 | 0.000 ± 0.009 | 0.5 ± 6000 | | 21 ± 1000 | 21 ± 353 | | *1* | *1* | | 1.035 ± 0.004 |
| | | 200ms | 610 | 0.00 ± 0.01 | 0.6 ± 8000 | | 24 ± 1000 | 24 ± 2000 | | *1* | *1* | | 1.033 ± 0.005 |
| | | 800ms | 610 | 0.00 ± 0.01 | 0.7 ± 6000 | | 24 ± 1000 | 24 ± 3000 | | *1* | *1* | | 1.020 ± 0.005 |
| | | 5s | 2079 | 0.000 ± 0.006 | 0.5 ± 6000 | | 12 ± 4000 | 12 ± 3000 | | *1* | *1* | | 1.02 ± 0.02 |
| S1D | H.-H. 2 component | 50ms | 44 | 0.000 ± 0.006 | 0.79 ± 3000 | | 43 ± 2000 | 42 ± 6000 | | *2* | *2* | | 1.022 ± 0.004 |
| | | 200ms | 36 | 0.000 ± 0.008 | 0.50 ± 8000 | | 44 ± 8000 | 45 ± 7000 | | *2* | *2* | | 1.022 ± 0.004 |
| | | 800ms | 42 | 0.000 ± 0.009 | 1 ± 5 | | 43 ± 38 | 30 ± 4000 | | *2* | *2* | | 1.006 ± 0.005 |
| | | 5s | 154 | 0.000 ± 0.005 | 0.6 ± 6000 | | 34 ± 4000 | 33 ± 5000 | | *2* | *2* | | 0.984 ± 0.008 |
| S1E | H.-H. 2 component | 50ms | 9 | 0.01 ± 0.005 | 0.9 ± 0.2 | | 61 ± 4 | 20 ± 20 | | *3* | *2* | | 1.02 ± 0.004 |
| | | 200ms | 4 | 0.005 ± 0.006 | 0.96 ± 0.03 | | 61 ± 3 | 10 ± 5 | | *3* | *2* | | 1.025 ± 0.006 |
| | | 800ms | 8 | 0.005 ± 0.006 | 0.91 ± 0.07 | | 63 ± 4 | 15 ± 10 | | *3* | *2* | | 1.007 ± 0.004 |
| | | 5s | 14 | 0.0002 ± 0.004 | 0.84 ± 0.07 | | 55 ± 3 | 14 ± 6 | | *3* | *2* | | 0.978 ± 0.007 |
| S1F | H.-H. 3 component | 50ms | 39 | 0.017 ± 0.007 | 0.79 ± 0.19 | 0.2 ± 530 | 75 ± 5 | 30 ± 30000 | 60 ± 2000 | *4* | *2* | *2* | |
| | | 200ms | 34 | 0.01 ± 0.01 | 0.78 ± 0.25 | 0.1 ± 440 | 75 ± 13 | 33 ± 4000 | 31 ± 5000 | *4* | *2* | *2* | |
| | | 800ms | 6 | 0.016 ± 0.006 | 0.85 ± 0.08 | 0.16 ± 0.07 | 76 ± 4 | 16 ± 10 | 3 ± 3 | *4* | *2* | *2* | |
| | | 5s | 7 | 0.008 ± 0.005 | 0.69 ± 0.09 | 0.28 ± 0.09 | 71 ± 4 | 20 ± 6 | 1 ± 2 | *4* | *2* | *2* | |
| 5 | H.-H. 2 component | 50ms | 6 | 0.015 ± 0.006 | 0.7 ± 0.2 | | 76 ± 4 | 30 ± 8 | | *4* | *2* | | 1.020 ± 0.004 |
| | | 200ms | 3 | 0.016 ± 0.005 | 0.91 ± 0.04 | | 73 ± 3 | 13 ± 5 | | *4* | *2* | | 1.024 ± 0.005 |
| | | 800ms | 7 | 0.015 ± 0.007 | 0.8 ± 0.1 | | 77 ± 5 | 22 ± 9 | | *4* | *2* | | 1.007 ± 0.005 |
| | | 5s | 7 | 0.008 ± 0.004 | 0.70 ± 0.08 | | 71 ± 4 | 19 ± 4 | | *4* | *2* | | 0.978 ± 0.007 |

Parameters are shown ± their standard deviation from the fit in IGOR Pro. Parameters in red are not properly determined, and those italicised were fixed. The goodness of fit ($\chi^2$) is coloured from worst (red) to best (green). The two component H.-H. fit with exponents $m_1$ and $m_2$ fixed to 1 is equivalent to a double exponential fit. The 1 and 2 component H.-H. functions are given in the methods, main text . The H.-H. 3 Component function (maximum fixed to 1) is as follows:

$$f(t) = y_0 + a_1 \cdot (1 - exp(-k_1 t))^{m_1} + a_2 \cdot (1 - exp(-k_2 t))^{m_2} + (1 - a_1 - a_2 - y_0) \cdot (1 - exp(-k_3 t))^{m_3}$$

### Supplementary Table 2 Sigmoid and Hill type fits to GluA2 recovery from desensitization

| Figure | Function | Duration of conditioning pulse | Goodness of fit ($\chi^2$) | Parameters | | | | |
|---|---|---|---|---|---|---|---|---|
| | | | | $y_0$ | $\tau$ | $m$ | $y_{max}$ | $t_{50}$ |
| S2A | Sigmoid | 50ms | 100 | 0.00 ± 0.01 | 0.0083 ± 0.0006 | | 1.01 ± 0.01 | 0.0314 ± 0.0008 |
| | | 200ms | 104 | 0.00 ± 0.02 | 0.0081 ± 0.0007 | | 1.01 ± 0.02 | 0.0305 ± 0.0009 |
| | | 800ms | 111 | 0.00 ± 0.02 | 0.0089 ± 0.0007 | | 0.99 ± 0.02 | 0.0313 ± 0.0009 |
| | | 5s | 157 | 0.00 ± 0.01 | 0.0095 ± 0.0006 | | 0.95 ± 0.01 | 0.0381 ± 0.0009 |
| S2B | Hill | 50ms | 9 | 0.014 ± 0.004 | | 2.5 ± 0.1 | 1.022 ± 0.004 | 0.0269 ± 0.0007 |
| | | 200ms | 5 | 0.009 ± 0.007 | | 2.5 ± 0.1 | 1.022 ± 0.004 | 0.0265 ± 0.0009 |
| | | 800ms | 8 | 0.008 ± 0.008 | | 2.4 ± 0.1 | 1.007 ± 0.004 | 0.0265 ± 0.0008 |
| | | 5s | 13 | 0.002 ± 0.005 | | 2.4 ± 0.1 | 0.975 ± 0.006 | 0.0317 ± 0.0008 |

Parameters are shown ± their standard deviations from fits in IGOR Pro. The parameter $y_0$ was constrained to be greater than zero for the sigmoid fits. The goodness of fit ($\chi^2$) is coloured from worst (red) to best (green) according to the scale in Supplementary Table 1.

Sigmoid function:

$$f(t) = y_0 + \left( y_{max} \middle/ \left( 1 + exp\left( \frac{t_{50} - t}{\tau} \right) \right) \right)$$

Hill equation:

$$f(t) = y_0 + (y_{max} - y_0) \,/\, \left( 1 + \left( \frac{t_{50}}{t} \right)^m \right)$$

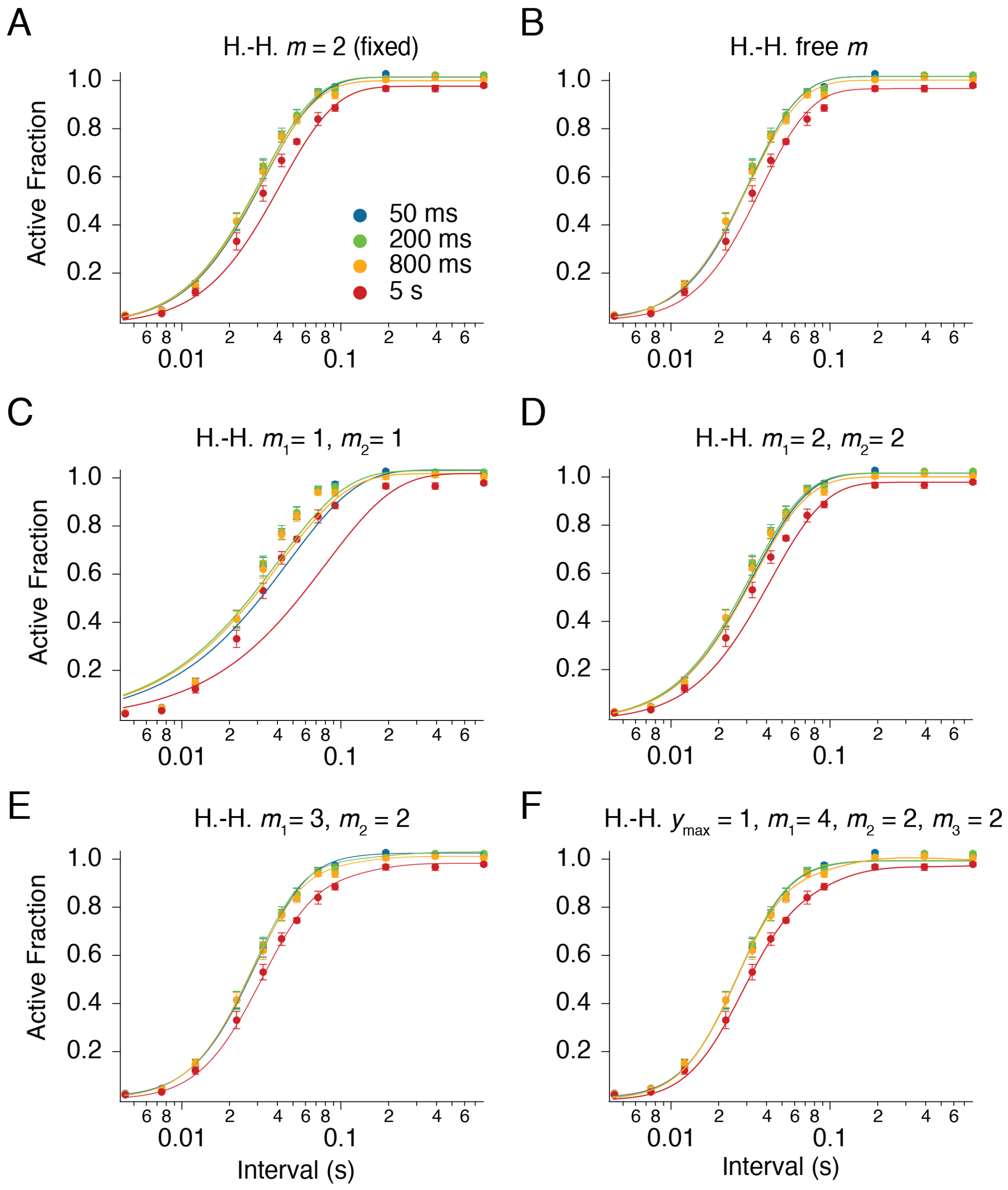


**Supplementary Figure 1 Hodgkin-Huxley fits to GluA2 recovery from desensitization**

Parameters and $\chi^2$ values for each fitted curve are given in Supplementary Table 1. Data points from Figure 5.

(A) Single component H.-H. fit with slope fixed to 2 cannot describe the recovery following a 5 s pulse (red). The early phase of recovery is poorly captured by this function because it is not steep enough.

(B) Allowing the slope to vary in a single component H.-H. function makes a better description of the early phase, but not for the recovery following the 5 s conditioning pulse (red curve). The disadvantage of a free slope is that rates and slopes are strongly negatively correlated during the fitting process.

(C) A two component H.-H. function with slopes fixed to 1 (equivalent to a double exponential) is a very poor descriptor, because the slopes are too shallow. Parameters in this fit are undetermined.

(D) A two component H.-H. function with both slopes fixed to two gives a poor description of the data following a 5 s conditioning pulse. For the shorter pulse data, the parameters are not determined.

(E) With one slope fixed to 3 and one slope fixed to 2, the description is good and all parameters are well determined. The goodness of fit ($\chi^2$) for each curve was similar to fits with $m_1 = 4$ and $m_2 = 2$ (Figure 5), except for recovery following the 5 s pulse, where the fit is definitively worse ($\chi^2 = 14$ vs. 7).

(F) The H.-H. function with three components (with fixed slopes, and maximum fixed to 1) gives a good description for recovery following the 5 s condition pulse. For shorter conditioning pulses, several parameters are not defined. For the recovery following the 800 ms pulse (orange curve), the slowest component ($k_3 = 3\ s^{-1}$) has a negative amplitude.

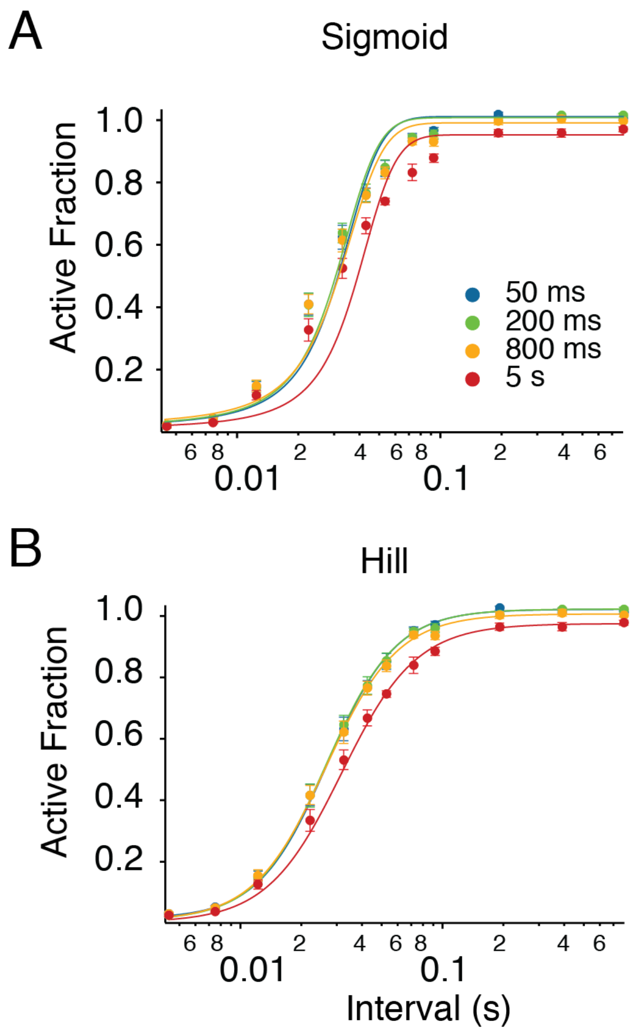


**Supplementary Figure 2 Sigmoid and Hill fits to GluA2 recovery from desensitization**
Equations, and parameters and $\chi^2$ values for each fitted curve are given in Supplementary Table 2 and its legend.
(A) Fits of sigmoid curves are extremely poor at describing the recovery, because they are too steep. The $y_0$ parameter was fixed to be positive.
(B) Allowing a free slope in the Hill equation describes the recovery from desensitization data surprisingly well, almost as well as any other function. The slope of the early phase of recovery is poorly described for the recovery after the 5 s conditioning pulse (red curve). The disadvantage of a free slope is that rate and slope are strongly negatively correlated during the fitting process. Overall, fitting the data with the Hill relation is problematic because the physical meaning of the equation with relation to conformational changes over time is unclear.
